# Structural and functional changes of pyramidal neurons at the site of an implanted microelectrode array in rat primary motor cortex

**DOI:** 10.1101/2022.09.15.507997

**Authors:** Bronson A. Gregory, Cort H. Thompson, Joseph W. Salatino, Mia J. Railing, Ariana F. Zimmerman, Bhavna Gupta, Kathleen Williams, Joseph A. Beatty, Charles L. Cox, Erin K. Purcell

## Abstract

Devices capable of recording or stimulating neuronal signals have created new opportunities to understand normal physiology and treat sources of pathology in the brain. However, it is possible that the initial surgical insertion and subsequent tissue response to implanted electrodes may influence the nature of the signals detected or stimulated. In this study, we characterized structural and functional changes in pyramidal neurons surrounding silicon or polyimide-based electrodes implanted in the motor cortex of rats. Devices were captured in 300 μm-thick tissue slices collected at the 1 or 6 week time point post-implantation, and individual neurons were assessed using a combination of whole-cell electrophysiology and 2-photon imaging. We observed disruption of the dendritic arbor of neurons near (<100 μm) the device surface at both time points, as well as a significant reduction in spine densities. These effects were accompanied by a decrease in the frequency of spontaneous excitatory post-synaptic currents (sEPSCs), a loss in sag amplitude, and an increase in spike frequency adaptation at the 6 week time point. Interestingly, we also noted a significant increase in filopodial density in neurons surrounding devices. Results were similar for polyimide and silicon-based electrodes. We hypothesize that the effects observed in this study may contribute to the signal loss and instability that often accompany chronically implanted electrodes.

## INTRODUCTION

Electrodes implanted in the brain enable the detection and stimulation of electrical and chemical signals from local neurons. Recent years have seen rapid growth in the design and application of neurotechnology in research and clinical settings^1–6^. New advances in the field have been driven by the movement to design softer, smaller, higher-density arrays with minimized insertional trauma^1,7^. Spikes from thousands of individual neurons can be simultaneously recorded from implants in rodents using state-of-the-art technology^8^, and hundreds of units have been isolated from arrays implanted in human subjects^9^. Ideally, implanted electrodes would serve as a “silent observer” of localized activity, allowing stable signal detection over many years without initiating a loss or functional impairment of signal-generating neurons. However, reductions in signal quality over time are often observed, and an expanding set of reported observations indicates substantial changes in the density, morphology, and gene expression of cells surrounding electrodes following implantation^10^.

The degree and time course of signal quality loss vary widely across literature, with some reports indicating relative stability, and others reporting pronounced and rapid losses. Chestek and colleagues^11^ reported an average 2.4% per month decline in spike amplitude recorded in non-human primates over variable observation periods (9-31.7 months), consistent with the observation that devices often lose activity over 6-12 months in non-human primates^12^. Comparatively more rapid decay in signal quality has been reported in the days and weeks following implantation in rodents^13^. In addition to chronic effects, signal instability has been reported within the span of a single day: in a study of acute recordings collected in human subjects, the majority of detected units displayed significant changes in their firing rate and spike amplitudes within a given recording session^14^. Similarly, data collected following several months of implantation in a non-human primate revealed substantial changes in spike amplitude within the span of a 1-hour recording session^15^. Non-stationarity introduces added complexity for systems that use decoding algorithms which rely on accurate spike detection, studies which interpret changes in firing rate to draw conclusions, and closed-loop strategies which deliver stimulation conditioned on spike detection. Various underlying cause(s) of these effects have been proposed, including natural physiological variability, mechanical/electrical sources of electrode failure, and the tissue response to electrodes post-insertion^10^,^16–18^.

Although the tissue response to implanted electrodes has long been hypothesized to be a key contributor to signal loss over long periods of time, defining the underlying mechanisms driving these responses, the link to recording quality, and the relationship to electrode design features all remain active areas of inquiry^19,20^. Electrode implantation results in an immediate and traumatic disruption of the neuropil and vasculature, followed by fluid extravasation and cellular infiltration into the implant site. The subsequent encapsulation of the device by responding reactive glia, and a loss of neuronal density, are commonly observed and have been used as metrics to assess the tissue response to implanted electrodes^10^,^21^. Changes in the structure and function of neurons are known effects of other forms of traumatic brain injury, yet these potential impacts have been comparatively less well-characterized in studies of implanted electrodes. Focal injuries of neocortex are known to result in a reorganization of network connectivity, local upregulation of glutamatergic transporters, changes in the expression of repolarizing potassium currents, and an altered balance of neurotransmitter receptors^22^. These injuries can produce hyperexcitability and seizure activity^22^, or alternatively, hypoexcitability^23,24^. Likewise, pronounced loss of dendritic spines is a known consequence of cortical impact^25^, which can persist for months following the initial trauma. Welle and colleagues observed disruption of dendrites surrounding electrodes through *in vivo* two-photon imaging^26^, but the impact of devices on dendritic spine densities and morphologies has not been explored previously. Recent reports have indicated changes in the expression of ion channels and synaptic markers at the gene and protein level surrounding implanted electrodes^27,28^, but effects on the intrinsic excitability of surrounding neurons are unknown.

In this study, we explored the hypothesis that electrode implantation alters the structure and function of local neurons. To do this, we captured single-shank, Michigan-style microelectrode arrays (MEAs) within thick sections of live rat motor cortex tissue collected at 1- and 6-weeks post-implantation. Whole-cell electrophysiology was used to assess changes in passive properties, spiking characteristics, and afferent synaptic activity in neurons “near” (<100 microns) or “distant” (~500 microns) from devices within individual tissue slices. During recording sessions, cells were filled with a fluorescent dye to assess changes in dendritic architecture and spine densities via 2-photon imaging and morphometric analysis. We tested both polyimide and silicon-based devices to assess material-based effects on responses. We observed several effects, which were pronounced in the neurons immediately surrounding the implant (<100 microns): (1) fluorescence microscopy showed that neurons displayed reduced dendritic length and branching at 1- and 6-weeks post-implant, (2) whole-cell intracellular recordings revealed that neurons surrounding devices showed reduced sag amplitude, increased spike frequency adaptation, and reduced frequency of spontaneous excitatory post-synaptic currents (sEPSCs) at 6 weeks, and (3) assessments of spine morphology indicated a loss of larger, more mature spine types accompanied by an increase in filopodia. The results suggest an emergence of a hypoexcitable network surrounding implanted electrodes which could contribute to observations of signal loss, while reductions in sag amplitude and increases in adaptation could contribute to variability in firing rates.

## METHODS

### Electrodes and surgical implantation

Two styles of single shank, 16-channel microelectrode arrays (MEAs) were chosen for comparison, one silicon-based and one polyimide-based. Non-functional, A1×16-style silicon devices were purchased from a commercial vendor (3 mm shank length, 15 μm thickness, tapered width measuring 123 μm at maximum)(Neuronexus Technologies, Ann Arbor, MI). Polyimide devices were custom-fabricated and supplied courtesy of Dr. John Seymour (University of Texas Health Science Center) based on methods previously described^29^. The dimensions of the polyimide devices were size-matched to those of the silicon devices, with the exception of a reduced thickness (4.4 μm). Devices were gas-sterilized prior to use. Either silicon or polyimide devices were bilaterally implanted into primary motor cortex (M1) of ≥12 week old male Sprague-Dawley rats based on methods and stereotaxic coordinates previously described^27–28^. Prior to implantation, the silicon devices were attached to a dummy connector and polyimide devices were temporarily fixed to a size-matched silicon shuttle via sterile 20% sucrose solution in PBS applied to the dorsal edge of the interface between the shuttle and device. Following insertion, the silicon devices were released from the dummy connector by severing the connection point with surgical scissors. The polyimide devices were released from the silicon shuttle by liberal application of sterile saline and withdrawal of the shuttle. Each device was inserted perpendicular to the cortical layers to a depth of 1.8-2.0 mm from the cortical surface of M1. The dorsal end of the device was flush or slightly protruding from the brain surface, and surgical closure was achieved with a combination of packed gel-foam and sterile suture of the incision site. Sham insertion/stab injury was achieved by brief insertion and immediate withdrawal of a silicon-based device following the craniotomy. Methods for sham insertions otherwise followed the same surgical procedures as for chronically implanted devices. All experimental procedures were performed in accordance with the National Institutes of Health *Guide for the Care and Use of Laboratory Animals* and approved by the Michigan State University Institutional Animal Care and Use Committee.

### Brain slice preparation

Animals were sacrificed at 1- or 6-weeks post-surgery to prepare brain slices targeting the implanted or sham-inserted region of M1. Briefly, rats were deeply anesthetized with 3% isoflurane and pentobarbital (100-200 mg/kg), transcardially perfused with slicing solution, and decapitated. Brains were quickly removed and placed into chilled (<4 °C), oxygenated (95% O_2_/5% CO_2_) slicing solution containing (in mM): 2.5 KCl, 1.25 NaH_2_PO_4_, 10.0 MgSO_4_, 0.5 CaCl_2_, 26.0 NaHCO_3_, 11.0 glucose, and 234.0 sucrose. Coronal slices (300 μm thickness) surrounding the implanted device or sham were obtained using a vibrating slicer (Leica Biosystems). The slices were then hemi-sectioned, and transferred to a chamber for 30 min of incubation in heated (36 °C) and oxygenated (95% O_2_/5% CO_2_) artificial cerebral spinal fluid containing (in mM): 126.0 NaCl, 2.5 KCl, 1.25 NaH_2_PO_4_, 2.0 MgCl_2_, 2.0 CaCl_2_, 26.0 NaHCO_3_, and 10.0 glucose.

### Electrophysiology

Slices of the implanted region were transferred to a submersion-type recording chamber and super-fused (2.5 ml/min) with oxygenated physiological solution maintained at 32 °C. Recording pipettes were pulled from 1.5 mm outer diameter capillary tubing and had tip resistances of 3–6 MΩ when filled with solution containing (in mM): 117.0 K-gluconate, 13.0 KCl, 1.0 MgCl_2_, 0.07 CaCl_2_, 0.1 EGTA, 10.0 HEPES, 2.0 Na_2_-ATP, 0.4 Na-GTP, and 50 μM Alexa Fluor 594. The pH of this solution was adjusted to 7.3 and osmolarity was adjusted to 290 mOsm. The use of this intracellular solution resulted in an 8 mV junction potential that has been corrected for in all voltage measurements.

Whole-cell recordings were obtained from deep layer pyramidal neurons in primary motor cortex with the visual aid of a BX51WI fixed-stage microscope (Olympus) equipped with Dodt contrast optics. A low-power objective was used to identify specific cortical layers and a high-power water immersion objective was used to visualize individual neurons. Electrophysiological data were acquired using a MultiClamp 700B amplifier filtered at 4 kHz and digitized at 10 kHz using a Digidata 1440A digitizer in combination with pCLAMP 10 software (Molecular Devices, San Jose, CA). Voltage clamp recordings were limited to neurons that had a stable access resistance of <20 MΩ. Current and voltage protocols were generated using pCLAMP software, and data were digitized and stored on a computer for off-line analyses.

Intrinsic properties and action potential (AP) output were collected in current-clamp configuration from neurons at resting membrane potential. Synaptic activity was recorded in voltage-clamp configuration while holding at –58 mV to isolate excitatory postsynaptic currents. Input resistance was calculated from the linear slope of the voltage-current relationship obtained from applying a series of depolarizing and hyperpolarizing constant current pulses from rest (−80 to +80 pA, 40 pA increments, 1s duration). Sag amplitude was calculated from a current step protocol that produced an initial hyperpolarization to ~-100 mV (1 s duration), and calculating the difference in voltage between the initial hyperpolarization and membrane potential at the end of the current step. Rheobase is measured as current (5 pA increments, 1 s duration) necessary to reliably evoke an action potential from rest. This current is then used to repeatedly evoke AP discharge (20 iterations). Action potential characteristics (threshold, half-width, and max amplitude) are measured for each iteration and averaged for each neuron. Firing properties of neurons were analyzed from a current step protocol that depolarized neurons from rest (0 to 2400 pA, 80 pA increment, 1s duration). Firing rate is reported as average over 1s depolarizing step. Spike frequency adaptation was calculated as the ratio of the mean of last 2 interspike intervals to the mean of the 3^rd^ and 4^th^ interspike interval. This calculation avoided the doublet AP discharge characteristic of neocortical pyramidal neurons. The slopes of the frequency-intensity relationship were calculated from the linear portion of the graph featuring at least 4 points above 0 pA. All population data are expressed as mean ± standard deviation. Statistical analyses were performed using GraphPad Prism (DotMatics, San Diego, CA) and Mini Analysis (Synaptosoft, Fort Lee NJ).

### Imaging

Deep layer pyramidal neurons filled via recording pipette with Alexa Fluor 594 (50 μM; Molecular Probes, Eugene, OR) were imaged by laser excitation (820 nm) using a two-photon laser-scanning microscopy system (Ultima, Bruker, Madison, WI) coupled with a Ti:Sapphire laser (Mai Tai HP, MKS-Spectra-Physics, Milpitas, CA). After at least 15 min of filling, each neuron was measured for consistent fluorescent levels, then basal dendrites were identified, and the entire length was recorded in 50 μm sections as a Z-stack of images for analysis of dendritic spines. Neurons were then imaged as Z-stacks of 200 μm x 200 μm sections overlapping by 20% to reconstruct the neuron’s full morphology by stitching the stacks together using Image J (NIH).

Dendritic arbors were reconstructed using stitched Z-stacks. Using the Simple Neurite Tracer tool in Image J, the traced dendrite arbors are skeletonized, allowing the length and branching patterns to be measured by Sholl analysis. Basal dendritic spines were reconstructed using the 50 μm sections taken as a Z-stack. Using NeuronStudio^30^, first the dendrite sections were reconstructed without dendritic spines using 3D voxels fit to the fluorescence region, then spines were identified by human observation and double-checked by a second observer. Spine morphology subtypes were determined (including thin, mushroom, stubby and filopodia) based on the spine’s head to neck diameter ratio and spine head diameter to spine neck length ratio^30^. Analysis for spine density and subtypes was carried out using exported data from NeuronStudio and GraphPad Prism (DotMatics, San Diego, CA).

### Statistics

Unless otherwise indicated, results are reported as Mean ± SD, and significance was determined at the P <0.05 level using a non-parametric, one-way ANOVA Kruskal-Wallis test, followed by Dunn’s test for multiple comparisons.

## RESULTS

### Dendritic Arborization is Altered in Near-Device Neurons

Visual inspection revealed an asymmetric distribution of dendritic arborization in Near-device neurons (defined as neurons within 100 μm of the device, Fig. 1A-B). Dendrites were preferentially lost on the implant-facing side of the cell, likely due to some combination of initial insertional injury and the subsequent chronic tissue response (Supp. Figs. 1-2). Quantification of these effects at the 1-week time point using Sholl analysis revealed reduced arborization (Fig. 1C) accompanied by a statistically significant loss in basal dendrite length in both Near-Silicon and Near-Polyimide neurons in comparison to distant neurons for both materials, as well as neurons in naïve, unimplanted tissue (Fig. 1D). At 6 weeks, Near-Silicon and Near-Polyimide neurons maintained the loss of basal dendrite length in comparison to naïve tissue (Fig. 1D). However, differences between Near-device and Distant-device cells (defined as neurons located ~500 μm from the device) no longer reached significance, which may be attributable to a slight decrease in the basal dendrite length of Distant-device neurons at the 6-week time point. Neurons near and distant from the insertional stab injury site in sham treatments displayed no significant difference from any other condition at either time point, indicating that the presence of the device was necessary to produce a significant loss of dendritic length. Likewise, there were no differences based on device material: both silicon- and polyimide-based electrodes produced similar effects.

**Fig. 1.**
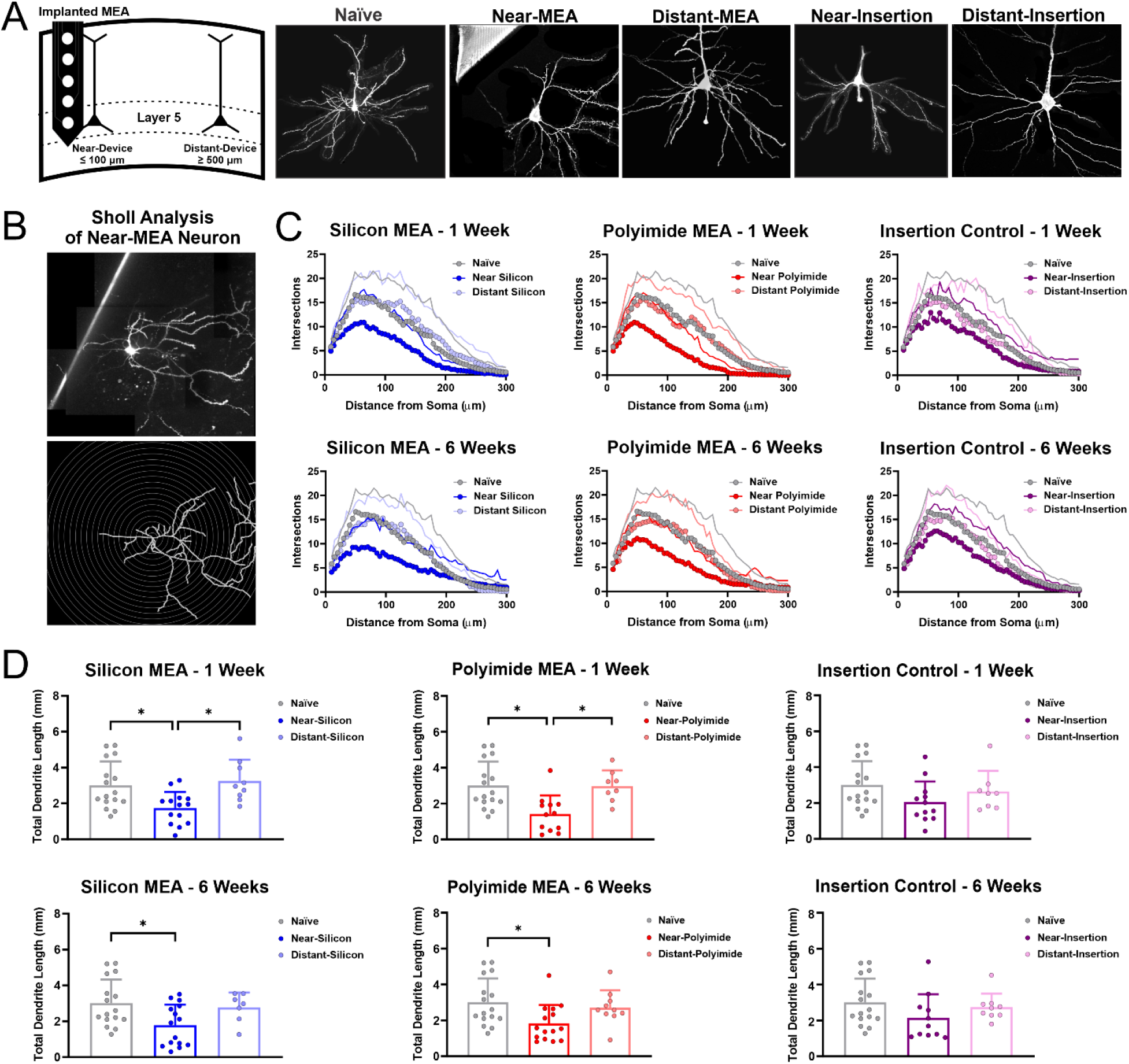
Electrode implantation is associated with disruption of the dendritic arbor in nearby neurons. (A) Schematic of implantation scheme and representative images of layer 5 regular-spiking pyramidal neurons in naïve tissue, as well as soma located near (<100 μm) or distant (~500 μm) from implanted electrodes or sham insertion sites (electrode can seen in top left corner of Near-MEA image). (B) Qualitative observation revealed an asymmetric loss of dendrites surrounding implants (top panel), which was quantified using Sholl analysis (skeletonized image of arbor, bottom panel). (C) Near-device neurons show a clear reduction in dendritic branching, as indicated by a reduction of intersecting points detected in Sholl analysis. (D) Overall dendritic length is significantly reduced in near-silicon and near-polyimide devices, which is not evident in sham insertion controls.

To further investigate electrode-based restructuring, we assessed the degree of dendritic branching in neurons surrounding implants. We found that Near-Silicon and Near-Polyimide neurons exhibited a reduction in dendritic branching, as indicated by decreased intersections with rings of increasing radii that are centered on the cell soma (Fig. 1B). For both polyimide and silicon-based electrodes, branching in Distant-device neurons was indistinguishable from naïve cells. Near-stab injury neurons displayed a comparatively mitigated loss of dendritic branching in comparison to Near-device neurons, reinforcing the impact of the indwelling device on observed disruptions to the dendritic arbor. Similarly to dendritic length, no material-based effects were observed in measurements of dendritic branching. Likewise, an asymmetric effect on branching was observed, where loss of branching was exacerbated on the implant-facing side of the cell (Supp. Fig. 3).

### Dendritic Spines are Less Dense and More Morphologically Immature in Near-Device Neurons

In addition to the loss of dendritic length and branching, we also noted a decrease in spine density (normalized to dendritic length) in Near-Device neurons. At the 1-week time point, spine density decreased by ~50% for both Near-Silicon and Near-Polyimide electrodes (Fig. 2A-B). Distant-Device neurons also displayed a significant, albeit somewhat lesser, reduction in spine densities for both electrode materials. While Near-Insertion neurons showed a ~30% reduction in spine density, no significant difference was observed between Distant-Insertion neurons and unimplanted controls. A modest improvement in spine density loss surrounding devices was observed by the 6-week time point, but reductions for Near-device, Distant-device, and Near-Insertion neurons remained significant in comparison to naïve controls (Fig. 2C). Spine losses for Near-Insertion neurons were relatively consistent between the 1- and 6-week time points in comparison to the slight recovery exhibited by neurons surrounding devices.

**Fig. 2.**
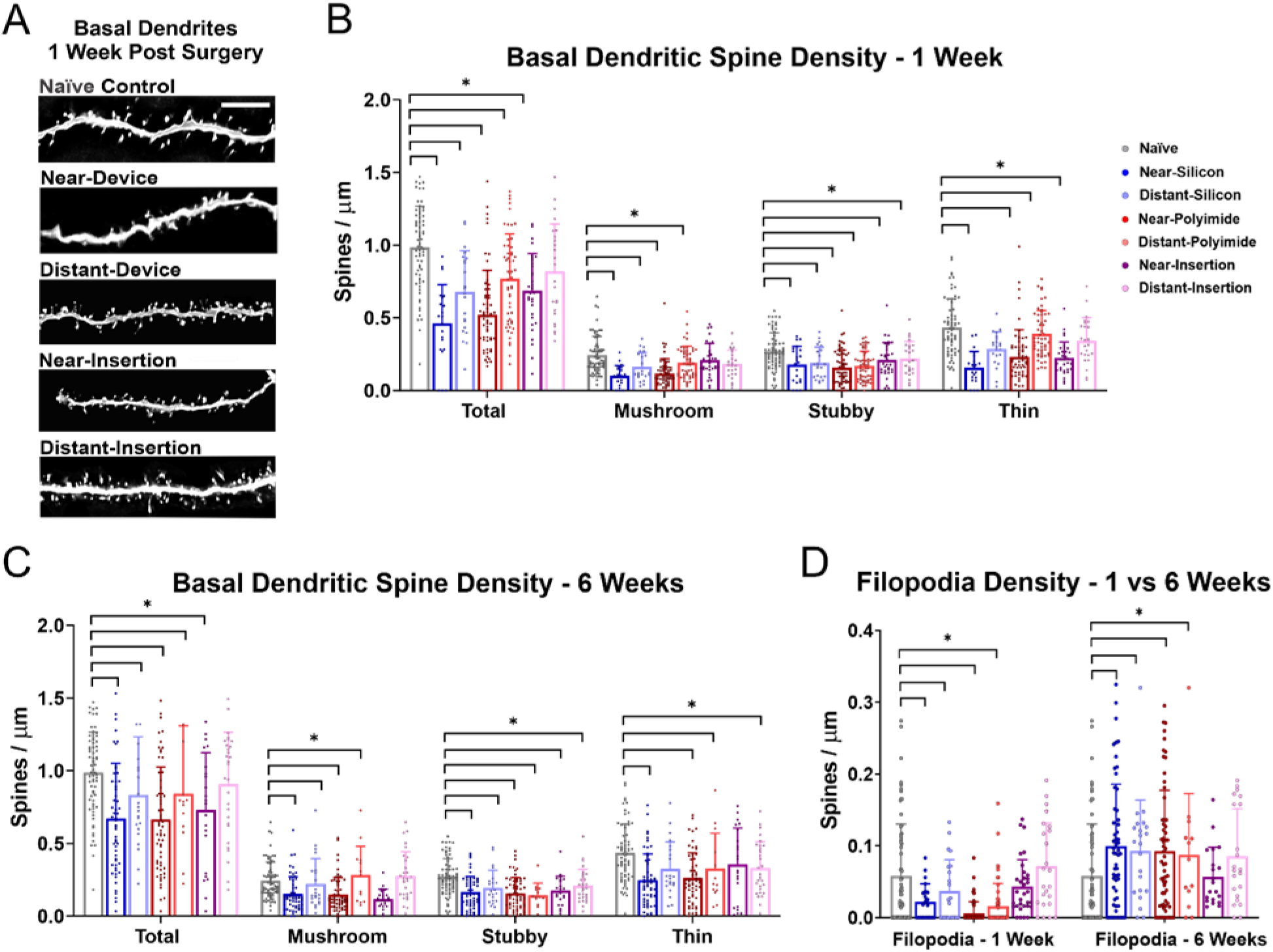
Dendritic spines are less dense and more morphologically immature surrounding devices. (A) Representative images of basal dendrites from Near-device, Distant-device, and Near-insertion pyramidal neurons after 1 week (scale bar = 10 μm). Near-device pyramidal neurons exhibited a lower total density of spines at both 1 Week (B) and 6 Weeks (C). Losses included reductions in spines with mushroom, stubby, and thin morphologies as well as filopodia. (D) Alternatively, the density of filopodia was increased surrounding devices at the 6-week time point. Filopodia density was not significantly affected in insertion injury controls.

We analyzed the morphology of individual spines and classified them according to common metrics reported in literature^31^. Mushroom spines, defined by a large head and a small neck, were significantly decreased only surrounding devices^32^. Stubby spines have similar functional characteristics but lack a well-defined neck. Stubby spines were significantly decreased at both time points in all conditions in comparison to naïve, unimplanted neurons. These losses were evident at both time points, irrespective of distance or electrode material. Thin spines are implicated in plasticity and are similarly shaped to mushroom spines, but with a smaller head. Spine losses were also observed in thin spines surrounding both devices and stab injury sites at each time point.

In contrast to the other spine types, filopodia were uniquely increased in density surrounding devices at the 6-week time point (Fig. 2D). Filopodia are long and thin, without a well-defined head. These structures are often observed in developing neurons, and glutamate uncaging studies indicate that they lack the AMPA receptors required for functional glutamatergic neurotransmission^33^. As such, they may mature into spines and are a putative site of “silent” synapses: synapses that are ultrastructurally normal, yet non-functional. Filopodia density was initially decreased surrounding devices in comparison to naive neurons at the 1-week time point, irrespective of distance or device material. However, this effect was reversed at the 6-week time point, when significant increases in filopodia density were observed. These effects were not recreated by the insertion injury. The emergence of filopodia surrounding devices could result from the loss in overall spine density at the 6-week time point.

### Near-Device Neurons Display Reduced Frequency of Spontaneous Excitatory Post-Synaptic Currents at 6 Weeks

Spontaneous excitatory post-synaptic currents (sEPSCs) are caused by neurotransmitter release from a presynaptic neuron in the absence of an applied stimulus. Initially, no detectable changes in the frequency or amplitude of sEPSCs were detected between any conditions at the 1-week time point (Fig. 3). At the 6-week time point, neurons near either silicon or polyimide-based electrodes displayed similar, significant reductions in the frequency of sEPSCs in comparison to naïve, unimplanted tissue. These effects required the presence of the device and were not recreated by the insertional injury. Similarly to the 1-week time point, this observation was decoupled from an effect on sEPSC amplitude at the 6-week time point, which could otherwise implicate changes in the activation of postsynaptic receptors.

**Fig. 3.**
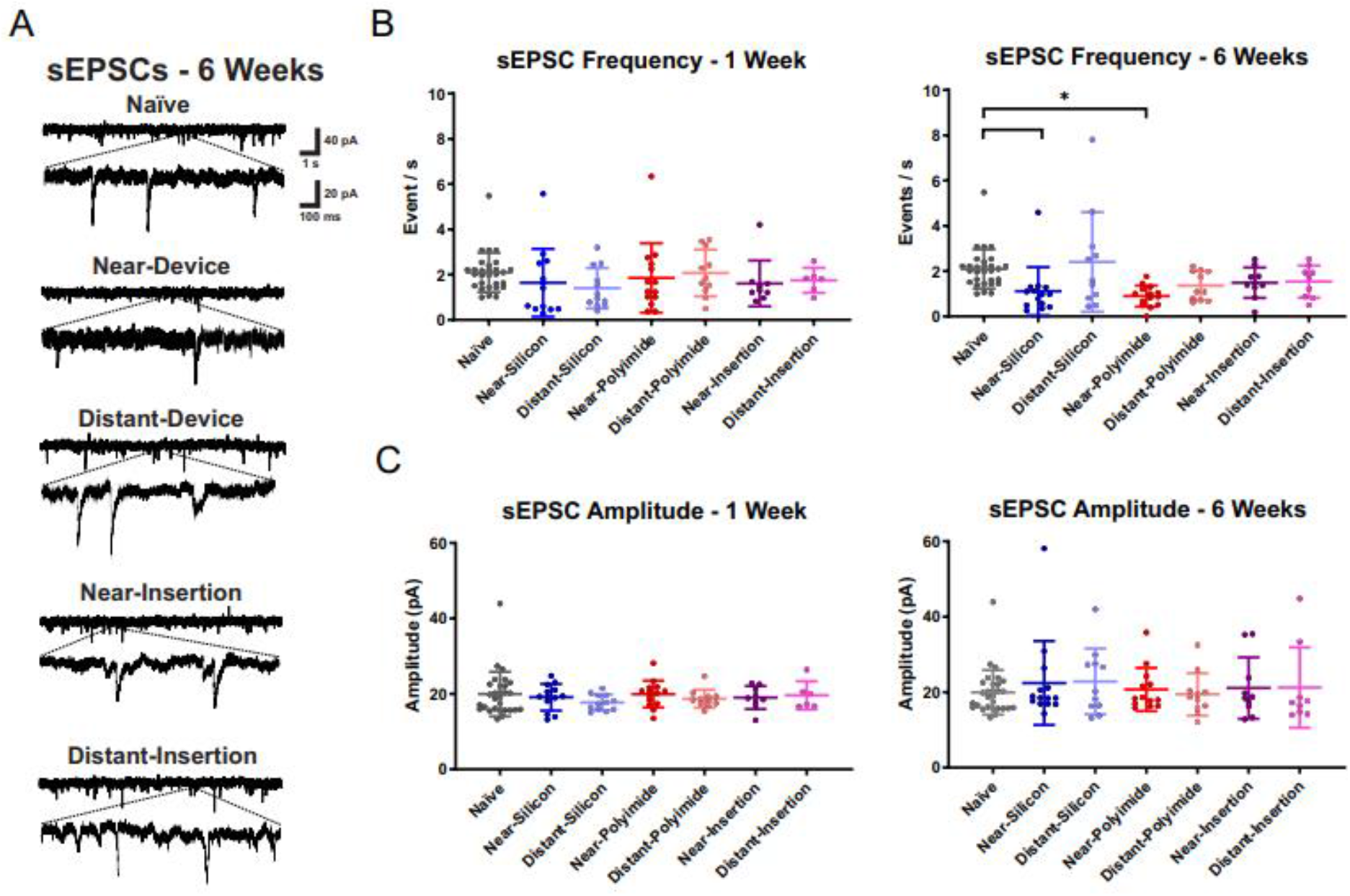
The frequency of spontaneous excitatory post-synaptic currents (sEPSCs) is reduced surrounding devices at 6 weeks. (A) Representative current traces from 6-week conditions. Spontaneous excitatory postsynaptic currents (sEPSCs) were measured with a Vhold of −58 mV. (B) The sEPSC frequencies were unaffected for Near-Device neurons at 1 week post-surgery, but at 6 weeks a significant decrease was observed only in neurons near devices (both polyimide and silicon). (C) sEPSC amplitudes appear unaltered in all conditions.

While structural changes in the dendritic arbor (Fig. 1-2) were accompanied by functional changes in sEPSC frequency in Near-Device neurons (Fig. 3), the timing of these effects was somewhat discordant. Significant losses in dendritic length, branching, and spine densities were observed at both the 1- and 6-week time points, but the reduction in sEPSC frequency was solely observed at the 6-week time point. The overall loss of spine density was the most pronounced at the 1-week time point, when no effects on sEPSC frequency were detected. Thus, although both structural and functional data indicate a generalized reduction in the underpinnings of synaptic transmission in Near-Device neurons, the observations implicate a contribution of alternative mechanisms in addition to losses in postsynaptic contact sites. Salatino et al. (2017) observed an initial increase in the expression of vesicular glutmate transporter 1 (VGLUT1) surrounding electrodes in rat primary motor cortex at three days post-implantation. Subsequent reduction of VGLUT1 and relatively increased expression of a GABA transporter (VGAT) were observed at the 4 week time point^27^. It is possible that increased VGLUT1 expression contributed to preserved excitatory tone near the device at the one week time point.

### Increased Adaptation and Reduced Sag Amplitude

Whole-cell electrophysiology was used to investigate the impacts of the implanted electrode on the intrinsic excitability of surrounding neurons. Measured passive and active properties included the resting potential, input resistance, rheobase, sag amplitude, spike frequency adaptation, firing frequency, the slope of the frequency-current relationship (FI Slope) and the characteristics of AP shape (Fig. 4). Complete results are summarized in Table 1. These measurements largely did not indicate any impacts of device implantation on neuronal intrinsic excitability, with two notable exceptions: the sag amplitude and spike frequency adaptation were both significantly affected at the 6-week time point in Near-Device neurons. Sag amplitude, which is a depolarization in response to a hyperpolarizing stimulus, was significantly decreased in neurons near both silicon and polyimide-based electrodes at the 6-week time point. These effects were not observed at the earlier time point or in stab injury controls. Increased spike frequency adaptation, which indicates broadening of the inter-spike interval during a sustained depolarizing stimulus, was observed in neurons near silicon and polyimide electrodes at the 6-week time point in comparison to naïve, unimplanted controls. Neurons near stab-wound insertion sites also displayed increased spike frequency adaptation at the 6-week time point, highlighting the importance of initial insertional damage in this effect. The reduction in sag amplitude, which controls the regularity of firing, as well as increased spike frequency adaptation, indicate that there are changes in the rhythmicity of neuronal firing immediately surrounding chronically implanted electrodes.

**Fig. 4.**
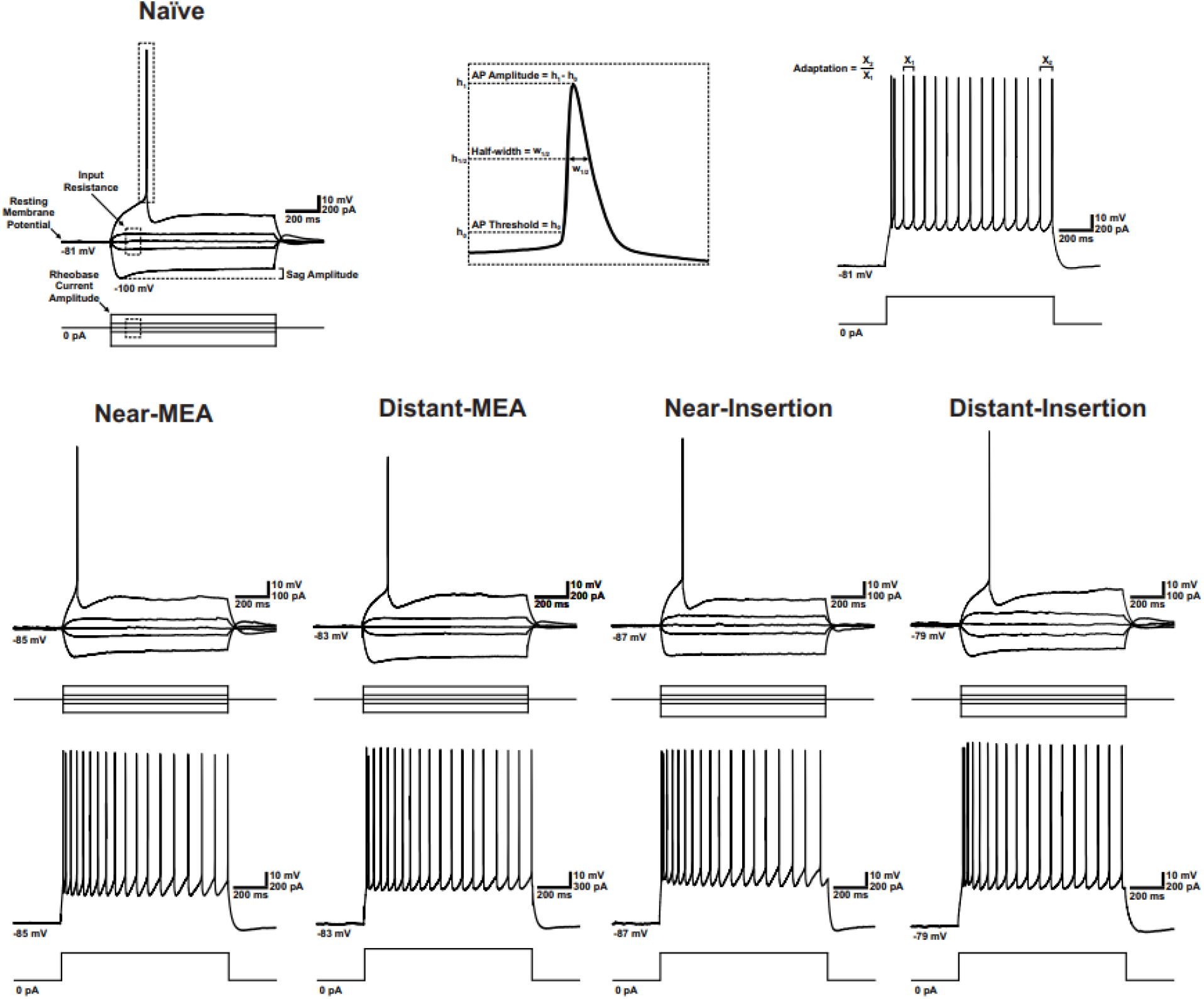
Assessing intrinsic excitability: an overview of electrophysiological characteristics analyzed. The features measured to test device-based impacts on active and passive electrophysiological properties are shown schematically (top panels), and representative images of firing responses are shown for neurons surrounding devices and insertion sites (bottom panels). Traces revealed qualitative effects of devices on spike frequency adaptation and sag amplitude. Quantitative data are summarized in Table 1.

**Table 1.** Summary of device-based impacts on electrophysiological properties of surrounding neurons. The majority of measurements showed no significant difference with neurons from naïve, unimplanted tissue. However, significant impacts of the device presence were observed at 6 weeks for the sag amplitude and the spike frequency adaptation: sag amplitude was significantly decreased, and adaptation was significantly increased versus controls (highlighted in gray, * P < 0.05).

|  |  | N | RMP<br>(mV) | R<sub>in</sub><br>(MΩ) | Sag<br>Amplitude<br>(mV) | Rheobase<br>(pA) | AP<br>Threshold<br>(mV) | AP Max<br>Amplitude<br>(mV) | AP Half-Width<br>(ms) | Max Firing<br>Frequency<br>(AP/s) | FI Slope<br>(AP/pA) | FI Adaptation<br>(Last/First IEI) |
| --- | --- | --- | --- | --- | --- | --- | --- | --- | --- | --- | --- | --- |
| Naïve | Mean | 44 | -78.0 | 99 | 5.9 | 165 | -45.0 | 84.5 | 0.72 | 49.6 | 0.056 | 1.49 |
|  | SD |  | ±5.68 | ±48.3 | ±3.65 | ±91.6 | ±5.39 | ±11.4 | ±0.162 | ±17.7 | ±0.0223 | ±0.477 |
| 1W Near-Silicon | Mean | 15 | -77.5 | 103 | 3.5 | 164 | -42.0 | 79.1 | 0.91 | 47.7 | 0.050 | 2.07 |
|  | SD |  | ±12.1 | ±50.3 | ±2.56 | ±113 | ±11.2 | ±17.2 | ±0.623 | ±21.6 | ±0.0201 | ±0.951 |
| 1W Distant-Silicon | Mean | 15 | -78.2 | 116 | 4.8 | 163 | -44.4 | 87.5 | 0.73 | 50.6 | 0.049 | 1.99 |
|  | SD |  | ±4.36 | ±41.8 | ±2.15 | ±107 | ±4.09 | ±5.75 | ±0.0934 | ±15.9 | ±0.0201 | ±0.738 |
| 1W Near-Polyimide | Mean | 23 | -74.5 | 109 | 5.4 | 164 | -40.9 | 79.4 | 0.78 | 32.7 | 0.052 | 1.61 |
|  | SD |  | ±15.3 | ±94.5 | ±4.09 | ±101 | ±8.30 | ±19.8 | ±0.244 | ±20.6 | ±0.0205 | ±0.577 |
| 1W Distant-Polyimide | Mean | 18 | -80.3 | 115 | 5.1 | 165 | -42.9 | 81.7 | 0.75 | 48.6 | 0.048 | 1.67 |
|  | SD |  | ±5.70 | ±52.3 | ±1.70 | ±99.9 | ±3.08 | ±10.1 | ±0.115 | ±17.8 | ±0.0151 | ±0.388 |
| 1W Near-Insertion | Mean | 15 | -72.9 | 112 | 6.2 | 161 | -42.6 | 81.8 | 0.81 | 42.6 | 0.054 | 1.71 |
|  | SD |  | ±10.3 | ±63.5 | ±4.67 | ±204 | ±3.42 | ±12.1 | ±0.151 | ±35.9 | ±0.0415 | ±0.965 |
| 1W Distant-Insertion | Mean | 13 | -78.5 | 104 | 5.5 | 176 | -42.0 | 88.4 | 0.67 | 59.6 | 0.070 | 1.43 |
|  | SD |  | ±4.22 | ±51.6 | ±3.18 | ±101 | ±10.6 | ±5.42 | ±0.0885 | ±17.8 | ±0.0355 | ±0.321 |
| 6W Near-Silicon | Mean | 18 | -74.0 | 119 | *2.9 | 150 | -45.8 | 91.6 | 0.75 | 60.0 | 0.058 | *2.340 |
|  | SD |  | ±11.7 | ±63.4 | ±2.27 | ±103 | ±4.93 | ±6.47 | ±0.0863 | ±8.32 | ±0.0246 | ±1.00 |
| 6W Distant-Silicon | Mean | 11 | -79.6 | 93 | 3.4 | 173 | -45.3 | 91.4 | 0.80 | 52.7 | 0.049 | 2.03 |
|  | SD |  | ±3.44 | ±33.3 | ±1.70 | ±58.7 | ±4.80 | ±8.24 | ±0.103 | ±13.2 | ±0.0131 | ±0.503 |
| 6W Near-Polyimide | Mean | 17 | -81.5 | 147 | *2.7 | 143 | -44.8 | 87.9 | 0.80 | 47.2 | 0.053 | *2.11 |
|  | SD |  | ±8.74 | ±96.9 | ±1.93 | ±90.3 | ±3.39 | ±7.25 | ±0.122 | ±17.6 | ±0.0299 | ±0.590 |
| 6W Distant-Polyimide | Mean | 12 | -80.3 | 93 | 5.5 | 208 | -46.0 | 82.5 | 0.75 | 44.3 | 0.040 | 1.89 |
|  | SD |  | ±6.46 | ±45.5 | ±3.33 | ±134 | ±4.88 | ±6.87 | ±0.104 | ±23.2 | ±0.0177 | ±0.944 |
| 6W Near-Insertion | Mean | 11 | -75.6 | 125 | 5.2 | 152 | -44.1 | 86.8 | 0.79 | 46.8 | 0.047 | *3.24 |
|  | SD |  | ±13.7 | ±62.0 | ±3.66 | ±105 | ±5.03 | ±7.91 | ±0.104 | ±21.1 | ±0.0299 | ±2.40 |
| 6W Distant-Insertion | Mean | 16 | -79.9 | 131 | 5.2 | 149 | -44.6 | 85.9 | 0.79 | 50.3 | 0.053 | 2.11 |
|  | SD |  | ±4.57 | ±81.6 | ±2.34 | ±89.6 | ±3.51 | ±6.79 | ±0.113 | ±20.9 | ±0.0164 | ±0.742 |

## DISCUSSION

While the biological response to electrodes implanted in the brain has long been viewed as a key contributor to signal loss and instability, a causal relationship between these phenomena has not been definitively established. Localized neuronal density loss and increased expression of glial markers are typically quantified to assess the degree of the tissue response to the electrode. However, these measurements are often misaligned in time and severity in comparison to the degree of signal loss, indicating that other factors are likely at play. While these effects may include electrical and mechanical failure points on the electrode, there is also a growing body of evidence that the biological response to electrodes is more complex than what can be captured by a few pre-selected markers^34–37^. The goal of our study was to add to current understanding of the impacts of electrodes on surrounding tissue, and our data illustrate nuanced structural and functional changes in the individual neurons within the recordable radius of the device interface^17^.

The seminal paper by Biran et al. (2005) reported a ~40% loss of neuronal density within the first 100 microns of a “Michigan”-style, single shank electrode implanted for a period of one month in the motor cortex of rats^21^. McConnell et al. (2009) attributed recording failure to an observed local “neurodegenerative state” marked by dendritic loss, neuronal death, and tau pathology within 100 microns of Michigan electrodes implanted in rat cortices^38^. Loss in neuronal density has since become a commonly used metric to assess the integration between implants and surrounding tissue. Less focus has been placed on the state of the remaining neurons. Using *in vivo* calcium imaging, Kozai and colleagues (2018) observed an initial reduction in neuronal activity surrounding implanted electrodes followed by functional recovery during the first month post-implantation^39^. Welle et al. (2020) showed progressive atrophy of the dendritic arbor surrounding electrodes during a three month observation period, likewise using *in vivo* 2-photon imaging^26^. Our methods are distinct from these previous studies in a few important ways: (1) our electrodes are implanted perpendicular to the cortical surface (rather than an oblique, shallow angle that facilitates *in vivo* imaging), (2) our morphological analysis was on a single cell basis, which allowed us to assess spine densities and perform Sholl analysis on individual arbors, and (3) our functional readouts are based on whole cell-electrophysiology, which allowed features of single action potentials, spike trains, and passive properties to be quantitatively characterized.

Similarly to previous reports, our data showed a generalized reduction in dendritic length surrounding implants. Our data also indicated that the loss of length, as well as reductions in branching, were asymmetrically driven by effects on the device-facing side. Additionally, our data provided new evidence of a reduction in spine densities surrounding electrodes implanted in the rat motor cortex (normalized to dendritic length). Since spines are typically the site of excitatory synapses, a loss in density implies a reduction in the network input to individual neurons surrounding electrodes. Loss in spine density was not restricted to the first 100 microns surrounding the device: it was also observed, albeit to a lesser extent, at the 500 micron distance. Reduced excitatory input to cells within 100 microns of the device could affect the spike output of individual neurons, posing a challenge to single unit detection. Loss of excitatory network input to cells at the 500 micron distance could influence the characteristics of the local field potential (LFP) in multiple ways. Perhaps the simplest interpretation would be that a generalized reduction in input could dampen the amplitude of the LFP due to reduced activity. However, the LFP is heavily influenced by the degree of neuronal synchrony and the geometric arrangement of the sources and sinks of ionic currents relative to the location of the electrode^40^. Generalized loss of excitatory input distally on dendrites could amplify the influence of inhibitory, somatic inputs on the frequency content of the LFP signal, potentially increasing relative power in higher frequency bands.

Asymmetry in the loss of the dendritic arbor, coupled with spine density loss, could influence the amplitude and frequency content of the LFP in unexpected ways^41–43^. Linden et al. (2011) reported a biophysical modeling approach to simulate LFP signals from activated, morphologically distinct cell populations, and found that the LFP amplitude for pyramidal neurons depends on correlated synaptic input, synapse location and neuron morphology^42^. Hence, the loss of dendritic arbor could possibly interfere with the conduction of synaptic input and diminish the spatial reach and amplitude of LFP. Another modeling study by Gold and colleagues (2006) observed that the extracellular action potential (EAP) waveform (defined by a wideband signal encompassing both the LFP and spike band) in CA1 pyramidal neurons was influenced by ion channel distributions more heavily than dendritic morphology^44^. Therefore, the impact of asymmetric dendritic loss on the LFP could result from the associated change in the spatial distribution of ion channels on individual neurons. More recently, Ness et al. (2016) used biophysical modeling to confirm that the LFP power generated from pyramidal neurons is heavily impacted by active subthreshold dendritic currents, and more importantly, by hyperpolarization activated currents (namely, the h-current)^45^. Assuming the loss of sag amplitude is reflective of a reduction in h-current, this study predicts a strong effect of electrode insertion on LFP power and frequency content. Predicting the impacts of our observations of neuronal restructuring on the LFP would benefit from dedicated modeling studies incorporating asymmetry and spine loss.

Morphological analysis suggested that remaining spines shifted toward a more immature, “silent” phenotype, as evidenced by an increase in filopodia surrounding electrodes. The increase in the density of filopodia can account for much of the recovery of spine densities between the 1- and 6-week time point. In glutamate uncaging studies, filopodia were associated with a lack of functional AMPA receptors, which were present on larger, stubby-type spines^33^. Absence of AMPA receptors would render these structures functionally silent, which would reinforce the generalized loss of excitatory input expected due to overall spine loss. In work published by Barres and colleagues (2005), ultrastructurally normal, but functionally silent, synapses have been associated with thrombospondin production by reactive astrocytes^46^. More recently, Liddelow et al. (2017) reported a reduction in excitatory synapses upon exposure to soluble cues from reactive astrocytes induced by microglial-derived inflammatory cytokines^47^. While the exact mechanisms are yet to be determined, we hypothesize that reactive glia contribute to the observed losses in neuronal spine densities, as well as the enrichment of filopodia. Using *in vivo* multiphoton imaging of mouse pyramidal neurons, Miyamoto et al. (2016) observed the formation of filopodia upon microglial contact with dendrites^48^. Additionally, Weinhard and colleagues (2018) found that microglia facilitate synaptic remodeling in mouse hippocampal cells by trogocytosis (i.e. selective, non-apoptotic partial phagocytosis^49^). Our recently reported gene expression data collected surrounding devices support the view that reactive glia are present at the device interface which are consistent with an inhospitable environment for synaptic transmission^35,36^.

We did not observe obvious effects of the device on spike shape, passive properties, or standard metrics of the responsiveness to stimulation. However, we did observe three significant effects of the device on the electrophysiological properties of local neurons. First, we observed reduced sag amplitude at the 6-week time point in neurons near polyimide and silicon devices. These effects were not present for Distant-Device neurons or stab insertion controls. To some extent, the reduction of sag amplitude may be related to generalized loss of the dendritic arbor, since the ion channels responsible for hyperpolarization-activated currents can be localized to the dendritic arbor. However, inspection of the morphological data indicates that there may be more to the story: while Near-Device neurons at the 1-week time point displayed marked dendritic loss, no significant change in sag amplitude was detected. This could suggest that a per-cell reduction in ion channel expression occurred in concert with dendritic loss; our previous data reported changes in ion channel expression surrounding devices^28^. Secondly, we observed increased adaptation in spike trains recorded in Near-Device neurons, as indicated by increasing interspike intervals in response to a constant stimulus. This effect may be related to the reduction in sag amplitude since hyperpolarization-activated currents can contribute to the rhythmicity of firing^50^. Finally, we observed a reduction in the frequency of spontaneous EPSCs in Near-Device neurons at the 6-week time point. While this may be partially attributable to spine loss, it is notable that sEPSC frequency is unaffected at the 1-week time point, when a more pronounced ~50% loss in spine density was observed. Again, a possible explanation involves a role for reactive glia: previous findings support that reactive microglia and astrocytes direct synaptic remodeling at device injury sites by releasing cytokines, glutamate, and adenosine triphosphate (ATP)^10,51,52^. Glial-derived adenosine produced by rapid hydrolysis of ATP can promote hyperpolarized post-synaptic contacts by downstream opening of K^+^ and Cl^-^ channels^10,53^. It is also possible that pre-synaptic mechanisms are also at play: we have reported a local, progressive reduction of VGLUT1 expression surrounding electrodes during the first month post-implantation^27^.

We did not observe any impacts of electrode material on our measured effects, despite a reduction in the Young’s modulus of polyimide in comparison to silicon. While several groups have pursued minimization of device-tissue modulus mismatch as a solution to poor integration, there are possible reasons for the lack of effect. First, recent reports implicate bending stiffness, which is a function of both Young’s modulus and device dimensions, as a stronger determinant of tissue response than Young’s modulus alone^54^. As such, the reduced Young’s modulus may have been insufficient to produce an impact on the tissue response without a coordinated reduction in dimensions. Secondly, the devices were pseudo-tethered, that is, stabilized only by surrounding gel foam and connective tissue infiltrating the craniotomy. We chose not to connectorize the implants in order to facilitate easy retrieval of devices within brain slices in this study, but it is possible that a more rigid fixation point would draw out the benefits of the polyimide material^55^. Investigating the impact of tethering forces will be a useful point of inquiry in future work.

In summary, we have reported a novel characterization of structural and functional changes in neurons surrounding implanted electrodes in the brain. While neurons present at the device interface retain the ability to produce signals, alterations in the regularity of spiking, the restructuring of neuronal arbors, and disengagement from the surrounding network could contribute to common observations of variability, signal loss, and shifting stimulation thresholds. The many opportunities for further investigation include extension of additional time points, materials, insertion strategies, and tethering schemes, as well as connecting functional read-outs to neuronal transcriptional profiles and underlying glial mechanisms. Future work is needed in this area to more completely characterize the effects of implanted electrodes on surrounding cells and predict effects of neuronal structural rearrangement on recorded signals via computational modeling.

## ACKNOWLEDGMENTS

The authors gratefully thank Dr. John Seymour (University of Texas Health Science Center, Houston, TX) for supplying the polyimide probes used in this study, and Sam Daniels for surgical assistance. This research was funded by NIH NINDS R01NS107451.

## Notes

### Competing Interest Statement

The authors have declared no competing interest.

