## Supplemental Figures 1-3 for "Structural and functional changes of pyramidal neurons at the site of an implanted microelectrode array in rat primary motor cortex"

### SUPPLEMENTARY FIGURES

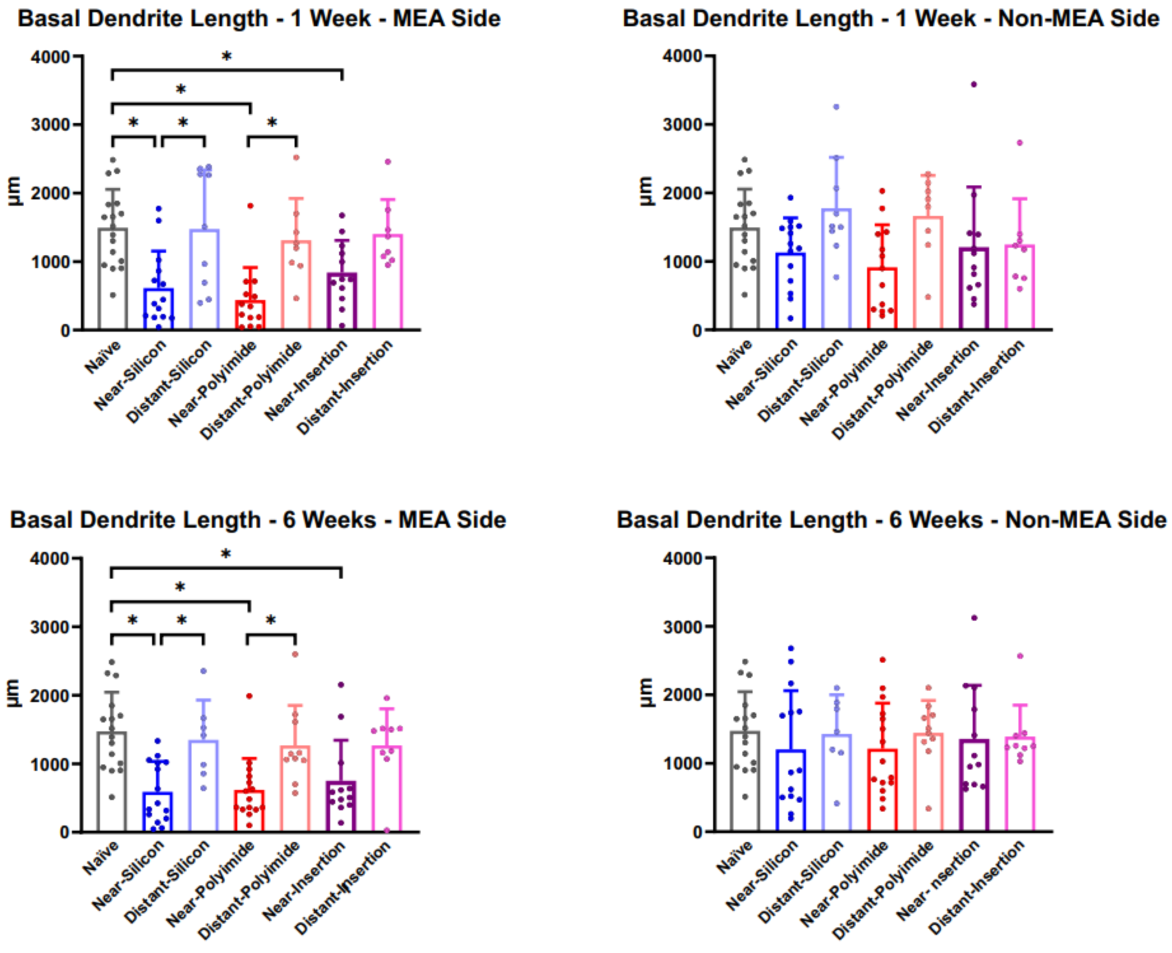


Supplementary Figure 1. Losses in dendritic length are asymmetric: reductions in overall length are driven by losses on the implant-facing side of the neuron.

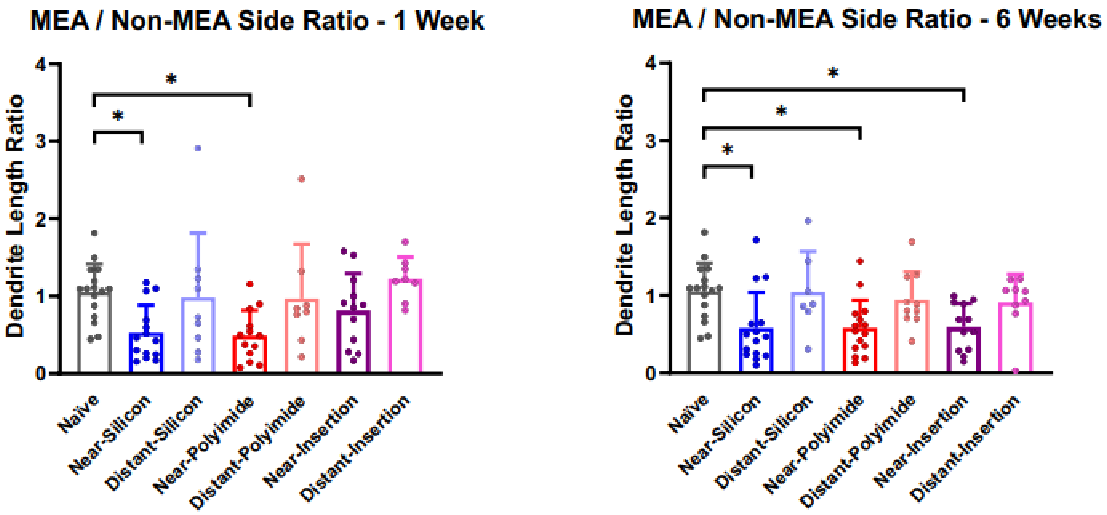


Supplementary Figure 2. Losses in dendritic length are asymmetric: the ratio of dendritic lengths from the implant-facing/non-facing sides reinforces the observation of preferential loss on the implant-facing side.

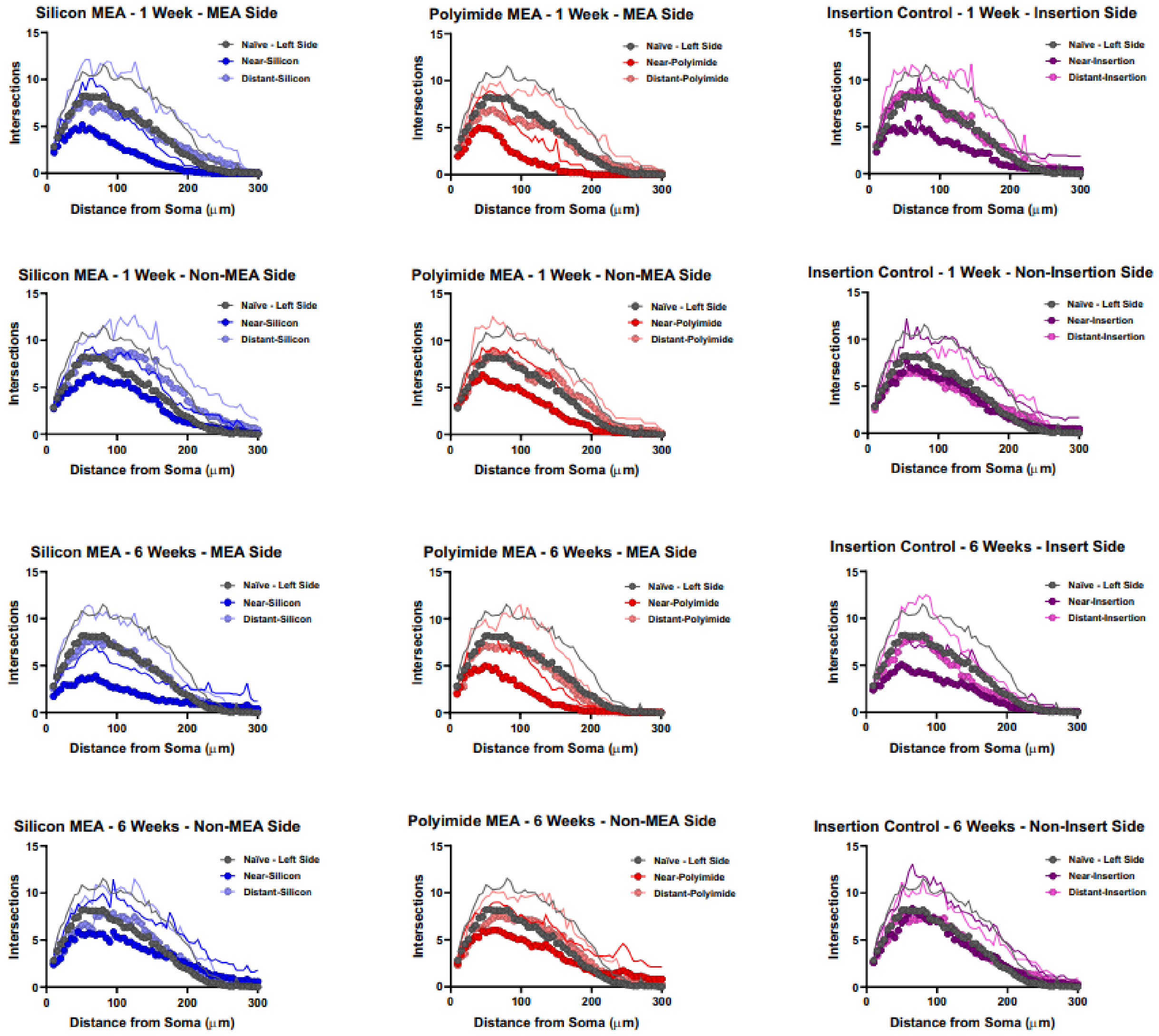


Supplementary Figure 3. Losses in dendritic branching are asymmetric: reductions in intersections detected in Sholl analysis are driven by losses on the implant-facing side.
